# A New Type of Satellite associated with Cassava Mosaic Begomoviruses

**DOI:** 10.1101/2021.03.11.434950

**Authors:** Catherine D. Aimone, Leandro De León, Mary M. Dallas, Joseph Ndunguru, José T. Ascencio-Ibáñez, Linda Hanley-Bowdoin

## Abstract

Cassava mosaic disease (CMD), which is caused by single-stranded DNA begomoviruses, severely limits cassava production across Africa. A previous study showed that CMD symptom severity and viral DNA accumulation increase in cassava in the presence of a DNA sequence designated as SEGS-2 (<u>s</u>equence <u>e</u>nhancing <u>g</u>eminivirus <u>s</u>ymptoms). We report here that when SEGS-2 is co-inoculated with *African cassava mosaic virus* (ACMV) onto *Arabidopsis thaliana*, viral symptoms increase. Transgenic *Arabidopsis* with an integrated copy of SEGS-2 inoculated with ACMV also display increased symptom severity and viral DNA levels. Moreover, SEGS-2 enables *Cabbage leaf curl virus* (CaLCuV) to infect a geminivirus resistant Arabidopsis accession. Although SEGS-2 is related to cassava genomic sequences, an earlier study showed that it occurs as episomes and is packaged into virions in CMD-infected cassava and viruliferous whiteflies. We identified SEGS-2 episomes in SEGS-2 transgenic Arabidopsis. The episomes occur as both double-stranded and single-stranded DNA, with the single-stranded form packaged into virions. In addition, SEGS-2 episomes replicate in tobacco protoplasts in the presence, but not the absence, of ACMV DNA-A. SEGS-2 episomes contain a SEGS-2 derived promoter and an open reading frame with the potential to encode a 75-amino acid protein. An ATG mutation at the beginning of the SEGS-2 coding region does not enhance ACMV infection in Arabidopsis. Together, the results established that SEGS-2 is a new type of begomovirus satellite that enhances viral disease through the action of a SEGS-2 encoded protein that may also be encoded in the cassava genome.

**IMPORTANCE:** Cassava is an important root crop in the developing world and a food and income crop for more than 300 million African farmers. Cassava is rising in global importance and trade as the demands for biofuels and commercial starch increase. More than half of the world’s cassava is produced in Africa, where it is primarily grown by smallholder farmers, many of whom are from the poorest villages. Although cassava can grow under high temperature, drought and poor soil conditions, its production is severely limited by viral diseases. Cassava mosaic disease (CMD) is one of the most important viral diseases of cassava and can cause up to 100% yield losses. We provide evidence that SEGS-2, which was originally isolated from cassava crops displaying severe and atypical CMD symptoms in Tanzanian fields, is a novel begomovirus satellite that can compromise the development of durable CMD resistance.

## INTRODUCTION

Cassava (*Manihot esculenta* Crantz), a major root crop in Africa, grows in acidic soil and under drought and high temperature conditions, making it an important staple crop for many smallholder farmers (FAO, 2010; Legg et al., 2015). Cassava mosaic disease (CMD) is one of the most devastating diseases of cassava (Nations and Africa, 2014). Losses due to CMD have an immediate impact on the food supply and threaten food security and the livelihoods of Africa’s rapidly growing population (FAO, 2010; Legg et al., 2015). CMD also causes serious problems for cassava production on the Indian subcontinent and increasingly in southeast Asia and China (Karthikeyan et al., 2016; Siriwan et al., 2020; Wang et al., 2019).

CMD is caused by a complex of at least 11 cassava mosaic begomoviruses (CMBs), of which nine occur in Africa and two are found on the Indian subcontinent (ICTV, 2019). They include *African cassava mosaic virus* (ACMV), *East African cassava mosaic virus* (EACMV), *East African cassava mosaic Cameroon virus* (EACMCV), *East African cassava mosaic Malawi virus* (EACMMV), *East African cassava mosaic Zanzibar virus* (EACMZV), *East African cassava mosaic Kenya virus* (EACMKV), *Cassava mosaic Madagascar virus* (CMMGV), *African cassava mosaic Burkina Faso virus* (ACMBFV), *Indian cassava mosaic virus* (ICMV), *South African cassava mosaic virus* (SACMV), and *Sri Lankan cassava mosaic virus* (SLCMV) (ICTV, 2019)

Begomoviruses constitute the largest genus of the *Geminiviridae*, a family of plant DNA viruses characterized by twin icosahedral particles (Stanley and Gay, 1983). They have small, circular single-stranded DNA (ssDNA) genomes that also occur as double-stranded DNA (dsDNA) replication intermediates in infected plants (Hanley-Bowdoin et al., 2000; Rojas et al., 2005).

Their genomes consist of either one or two DNA components ranging in size from 2.5-2.9 kb (Zerbini et al., 2017). CMBs have two genome components designated as DNA-A and DNA-B (Briddon et al., 2010). The DNA-A component can replicate autonomously and produce virions in infected plants, while DNA-B is required for cell-to-cell and systemic movement (for review see (Hanley-Bowdoin et al., 2013)). DNA-A encodes 5-6 proteins involved in replication, transcription, enscapsidation and countering host defenses while DNA-B encodes two proteins necessary for movement (Hanley-Bowdoin et al., 2013). Like all begomoviruses, CMBs are transmitted by whiteflies (*Bemisia tabaci* Genn) (Mugerwa et al., 2012). They are also spread by vegetative propagation of stem cuttings from CMD-infected cassava (FAOSTAT, 2016).

CMBs evolve rapidly via a combination of nucleotide mutation, recombination and genome reassortment (Duffy and Holmes, 2009; Pita et al., 2001; Zhou et al., 1997), which can result in the emergence of new and more virulent viruses (Lefeuvre and Moriones, 2015; Sanjuán et al., 2010). CMBs often occur in mixed infections leading to synergy between the different viruses and increased symptom severity (Fondong et al., 2000; Pita et al., 2001). In the 1990s and 2000s, synergy of a CMB recombinant between ACMV and EACMV contributed to a severe pandemic that spread from Uganda to other sub-Saharan countries and devastated cassava production (Deng et al., 1997). In response to the pandemic, many African farmers adopted cassava cultivars with the CMD2 locus, which confers resistance to CMBs (Akano et al., 2002; Rabbi et al., 2014).

Many begomoviruses are associated with satellite DNAs that are packaged into virions and can increase virulence and alter host range (Briddon et al., 2003; Briddon et al., 2018a; Mansoor et al., 2003). To date, 3 types of DNA satellites have been described, including alphasatellites, betasatellites, and deltasatellites (Hassan, 2016; Nawaz-ul-Rehman and Fauquet, 2009; Zhou, 2013). Alphasatellites and betasatellites are approximately 1300-1400 nt in size, contain one major open reading frame (ORF), a hairpin structure, and an adenine-rich region (Hassan, 2016; Nawaz-ul-Rehman and Fauquet, 2009; Zhou, 2013). Deltasatellites are related to betasatellites but range from ca. 540 nt to750 nt in size (Hassan, 2016). Betasatellites rely on the Rep protein of their helper virus for replication whereas alphasatellites encode their own Rep protein and replicate autonomously (Briddon et al., 2018b; Rosario et al., 2016).

Betasatellites encode a single protein, βC1, that enhances geminivirus symptoms by suppressing post-transcriptional gene silencing (PTGS) and transcriptional silencing (TGS) (Gnanasekaran et al., 2019; Yang et al., 2019). Some alphasatellites display silencing suppression activity and increased symptom severity (Nawaz-ul-Rehman and Fauquet, 2009), while others are associated with attenuation of symptoms (Idris et al., 2011). Deltasatellites are noncoding DNAs that can reduce the titer of associated begomoviruses (Hassan, 2016). An alphasatellite has been found in CMB-infected cassava in Madagascar off the coast of East Africa (Harimalala et al., 2013), but there are no reports of satellites associated with CMD on the African continent. However, coinoculation studies in *Nicotiana benthamiana* showed that some CMBs can interact with heterologous beta and alpha satellites to impact symptoms and viral DNA accumulation (Patil and Fauquet, 2010).

Two novel DNAs, designated SEGS-1 (DNA-II, AY836366) and SEGS-2 (DNA-III, AY836367; <u>s</u>equences <u>e</u>nhancing <u>g</u>eminivirus <u>s</u>ymptoms) were isolated from cassava plants showing severe, atypical CMD symptoms in fields near the Tanzanian coast (Ndunguru et al., 2016). SEGS-1 and SEGS-2 both contain regions with high GC content, but are distinct DNAs that only share 23% sequence identity. Sequences related to both SEGS occur in the cassava genome, with the most closely related copies displaying 99% and 84-87% identity to the cloned SEGS-1 and SEGS-2 sequences, respectively. Controlled inoculation experiments showed that both SEGS-1 and SEGS-2 enhance CMD symptoms in cassava (Ndunguru et al., 2016).

Both SEGS-1 and SEGS-2 occur as circular episomes in infected but not healthy cassava leaves. SEGS-2 episomes are also found in DNA from viruliferous whiteflies and in virions from cassava and whiteflies, indicating that SEGS-2 is packaged and acquired by whiteflies (Ndunguru et al., 2016). The SEGS-2 episome includes a 52-bp sequence that does not match the cassava genome but contains a 26-bp motif related to alphasatellite sequences. The alphasatellite motif may reflect a recombination event between the cassava genome and an alphasatellite (Ndunguru et al., 2016).

The presence of sequences related to SEGS-2 in the cassava genome has complicated functional studies of SEGS-2. To overcome this constraint, we examined SEGS-2 during CMB infection of *Arabidopsis thaliana* plants and *Nicotiana tabacum* suspension cells. These studies provide evidence that SEGS-2 is a new type of begomovirus satellite associated with disease enhancement.

## RESULTS

### SEGS-2 enhances ACMV symptoms in Arabidopsis

To establish *Arabidopsis* as a model system for studying SEGS-2, we first asked if ACMV can infect *Arabidopsis thaliana*. A previous report showed that the related CMB, SACMV, can infect the *Arabidopsis* Col-0 accession (Pierce and Rey, 2013), but we were unable to infect Col-0 plants with ACMV. Hence, we determined if the hypersusceptible *Arabidopsis* accession, Sei-0 (Lee et al., 1994), can be infected with ACMV when co-bombarded with partial tandem dimers of DNA-A and DNA-B (Fig. 1A-B). As shown in Fig. 1B, we observed ACMV infection in Sei-0. However, the timing of symptom appearance varied between experiments, most likely because ACMV is not well adapted to *Arabidopsis* as a host. For this reason, we assayed 9 plants/treatment and only compared treatments within an experiment in which the plants were grown and inoculated together. Conclusions are based on three replicates for each experiment.

**FIG 1.**
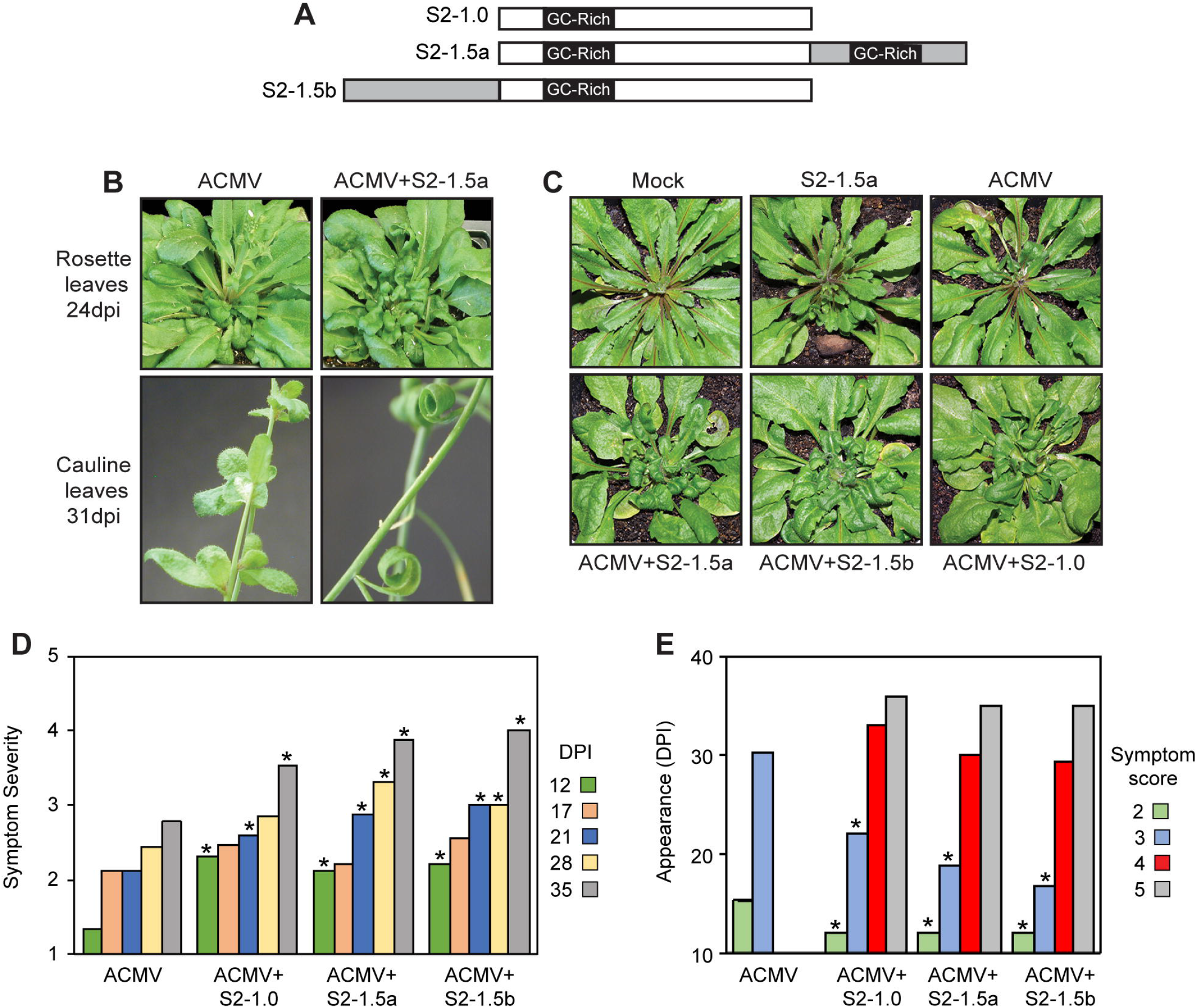
SEGS-2 enhances CMB infection in *Arabidopsis thaliana*. (A) Diagram of SEGS-2 clones used for infection studies. S2-1.0 contains a single copy of SEGS-2. S2-1.5a and S2-1.5b are partial tandem dimers with duplicated regions shown in grey. The GC-rich region is indicated in each construct. (B) Rosette and cauline leaves of wild-type Arabidopsis Sei-0 plants inoculated with ACMV DNA-A + DNA-B alone (left) or in combination with S2-1.5a (right). The rosette and cauline images are at 24 and 31 dpi, respectively. (C) Arabidopsis Sei-0 inoculated with ACMV alone or in combination with S2-1.0, S2-1.5a, or S2-1.5b at 25 dpi. (D) Symptom score on a scale of 1 [none] to 5 [strong chlorosis, stunting, and leaf deformation] for wild-type Sei-0 inoculated with ACMV alone or in combination with S2-1.0, S2-1.5a, or S2-1.5b from 12-35 dpi. (E) Average day post infection of symptom appearance for wild-type Sei-0 inoculated with ACMV alone or in combination with S2-1.0, S2-1.5a, or S2-1.5b.

To examine the impact of SEGS-2 on ACMV infection, we generated a set of SEGS-2 constructs (Fig. 1A) for co-bombardment with the infectious ACMV clones. The constructs were derived from the SEGS-2 clone originally amplified from CMB-infected cassava using an alphasatellite universal primer (Ndunguru et al., 2016). S2-1.5a and S2-1.5b are partial tandem dimers with different halves of SEGS-2 duplicated. The design of the SEGS-2 1.5-mer clones was based on begomovirus infectious clones. We also generated a SEGS-2 monomer clone (S2-1.0) that does not contain primer sequences that were introduced during construction of the original SEGS-2 clone.

Sei-0 plants were bombarded with ACMV DNA-A and DNA-B infectious clones alone or in the presence of a SEGS-2 clone. Plants inoculated only with ACMV showed very mild symptoms on rosette and cauline leaves at 24 and 31 dpi, respectively (Fig. 1B). In contrast, plants co-bombarded with ACMV and SEGS-2 displayed strong leaf curling and stunting at the same times (Fig. 1B). Severe symptoms were also apparent at 25 dpi on plants (Fig. 1C) co-inoculated with ACMV and one of the SEGS-2 clones in Fig. 1A (S2-1.5a, S2-1.5b, or S2-1.0). Symptoms were monitored over time (12 to 35 dpi) using a scale of 1 [none] to 5 [strong chlorosis, stunting, and leaf deformation] (Fig. 1D and 1E). The average symptom scores for plants inoculated with ACMV alone were consistently lower than those for plants inoculated with ACMV and SEGS-2 (Fig. 1D). The differences were statistically significant at 12, 21, 28, and 35 dpi in comparisons between ACMV alone and ACMV+SEGS-2 plants. The average time of symptom appearance was also later for plants inoculated with ACMV alone than for plants inoculated with ACMV+SEGS-2 (Fig. 1E). The delay was significant for symptom scores 2 and 3. Strikingly, symptom scores 4 and 5 were only observed in plants inoculated with ACMV + SEGS-2. There were no differences in the average symptom scores and time of symptom appearance for the different SEGS-2 clones. Together, these results showed that like cassava (Ndunguru et al., 2016), SEGS-2 enhances ACMV infection in Sei-0 plants, establishing that *Arabidopsis* can be used as a model system to study SEGS-2.

### A SEGS-2 transgene enhances ACMV symptoms

The cassava genome contains sequences related to SEGS-2, prompting us to ask if SEGS-2 can enhance ACMV infection when it is integrated into the *Arabidopsis* genome. We produced transgenic Sei-0 lines with the SEGS-2 monomer sequence cloned into the Ti plasmid DNA vector in both orientations. The transgenic plants were readily recovered and appeared phenotypically normal, indicating that the SEGS-2 sequence by itself does not have a visible impact on *Arabidopsis* plants.

SEGS-2 transgenic plants were inoculated with ACMV alone, and symptoms and viral DNA accumulation were examined. Transgenic plants carrying SEGS-2 in either orientation (T-S2-1.0F or T-S2-1.0R) displayed strong symptoms at 28 dpi, while wild-type Sei-0 plants only showed mild symptoms (Fig. 2A). The average symptom scores of the transgenic lines were significantly higher than wild-type plants at 14, 21 and 28 dpi but not at 35 dpi (Fig. 2B). Co-bombardment of the transgenic plants with S2-1.5b plasmid DNA + ACMV did not increase symptom severity (Fig. 2C), indicating that the SEGS-2 transgene was sufficient for maximal activity. The transgenic plants also accumulated viral DNA earlier and at higher levels than wild-type plants (Fig. 2D). Viral DNA was readily detected in the transgenic plants at 14 and 21 dpi by semi-quantitative PCR using ACMV DNA-A specific primers. In contrast, viral DNA was first detected in wild-type at 28 dpi, and viral DNA levels were considerably higher at 28 and 35 dpi in transgenic versus wild-type plants. Importantly, enhancement of symptoms and viral DNA accumulation was independent of the orientation of SEGS-2 in the Ti plasmid vector (cf. T-S2-1.0F and T-S2-1.0R), indicating that flanking T-DNA sequences did not contribute to disease enhancement.

**FIG 2.**
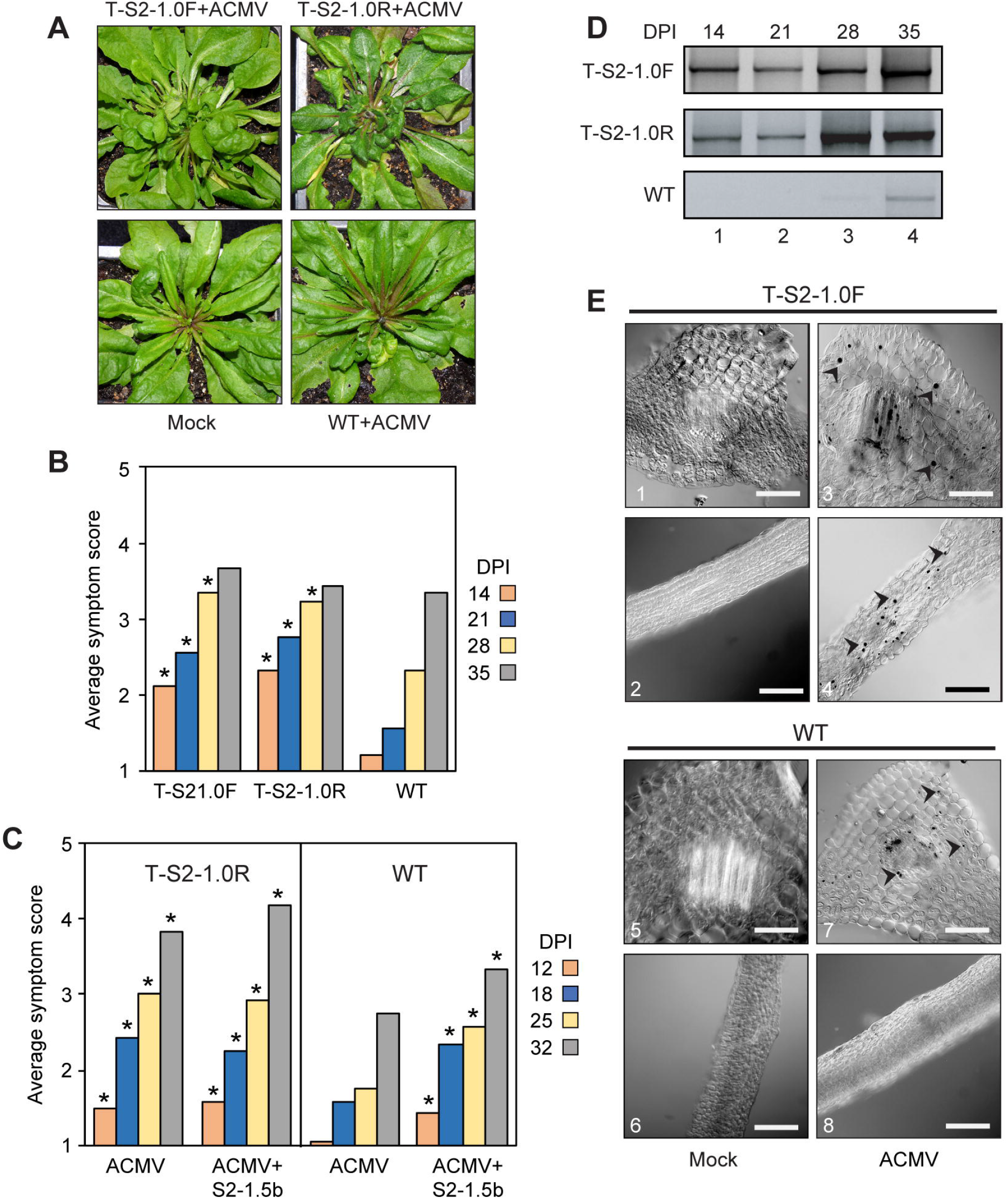
A SEGS-2 transgene enhances CMB infection in Arabidopsis. (A) Transgenic Sei-0 plants (T-S2-1.0F or T-S2-1.0R) inoculated with ACMV alone show severe symptoms at 28 dpi, while wild-type (WT) Sei-0 plants (WT) display mild symptoms. No symptoms were observed on plants inoculated with ACMV-B alone (Mock). (B) Average symptom scores for T-S2-1.0F, T-S2-1.0R plants and wild-type Sei-0 plants inoculated with ACMV at 14, 21, 28, and 35 dpi. *P < 0.05 from a Wilcoxon rank sum test of comparisons between SEGS-2 transgenic and wild-type plants. (C) Average symptom scores for T-S2-1.0F plants and wild-type (WT) plants inoculated with ACMV alone or with ACMV+S2-1.5b. *P < 0.05 from a Wilcoxon rank sum test of comparisons to wild-type plants (WT). (D) Total DNA was extracted at 14 (lane 1), 21 (lane 2), 28 (lane 3), and 35 (lane 4) dpi from T-S2-1.0F, T-S2-1.0R, and wild-type Sei-0 plants and analyzed by semi-quantitative PCR using the CMAfor2/CMArev2 primer pair for ACMV DNA-A. (E) Tissue sections of T-S2-1.0F (top) and wild-type (WT; bottom) Sei-0 plants at 35 dpi incubated with a digoxigenin DIG-labeled DNA probe for ACMV A and visualized using an anti-DIG antibody conjugated to AP-conjugated antibody. Arrowheads mark infected cells containing ACMV-A DNA in their nucleus (3, 4 and 7). No virus-positive cells were observed in mock-inoculated controls (1, 2, 5 and 6) or in wild-type leaf sections inoculated with ACMV (8). Bars represent 1 micron at 20X magnification.

We used *in situ* hybridization to examine the pattern of ACMV infection in the presence and absence of SEGS-2. These studies used an ACMV-A oligonucleotide probe conjugated to digoxigenin that specifically binds to viral DNA and an anti-digoxigenin detection system that stains virus-positive nuclei with a dark precipitate. Petiole sections from T-S2-1.0F plants did not contain more virus-positive nuclei than wild-type plants, but the staining appeared to be stronger in T-S2-1.0F petiole sections (Fig. 2E). Stained nuclei were also observed in the leaf sections from T-S2-1.0F plants but not from wild-type plants. Many of the stained nuclei in the T-S2-1.0F leaf sections were near vascular bundles but some were more dispersed. No staining was observed in leaf sections from the mock inoculated controls, establishing the specificity of the *in situ* assay. These results suggested that SEGS-2 enhances ACMV infection by increasing the target cell population in the host.

### Episomal copies of SEGS-2 in *Arabidopsis thaliana*

SEGS-2 episomes have been detected in infected cassava leaves and viruliferous whiteflies collected from cassava fields (Ndunguru et al., 2016). Hence, we asked if SEGS-2 episomes also occur in Arabidopsis. Total and virion DNA preparations were used as templates for rolling circle amplification (RCA), which preferentially amplifies small, circular DNA molecules. The RCA products were then subjected to PCR using divergent primer pairs, (2-4F/2-6R) that amplify SEGS-2 episomal DNA but not genomic sequences (Ndunguru et al., 2016) (Fig. 3A). SEGS-2 episomes were detected in ACMV-infected plants carrying the T-S2-1.0F transgene at 37 dpi (Fig. 3B; lane 4), but not in infected wild-type plants (lane 2). When the gel image was enhanced 10-fold, a weak band corresponding to the SEGS-2 episome was also visible in mock-inoculated transgenic plants (Fig. 3B; lower panel, lane 3) but not in equivalent wild-type plants (lane 1). We also detected SEGS-2 episomes in virion DNA from ACMV-infected T-S2-1.0F plants (Fig. 3C; lane 2) but not in samples prepared in parallel from mock-inoculated transgenic plants (lane 1). Contamination of the RCA products by *Arabidopsis* genomic DNA was ruled out using the PCNA2-F/R primer pair, which amplifies a 264-bp product corresponding to the host gene (AT2G29570) encoding proliferating cell nuclear antigen (Fig 3C; bottom panel).

**FIG 3.**
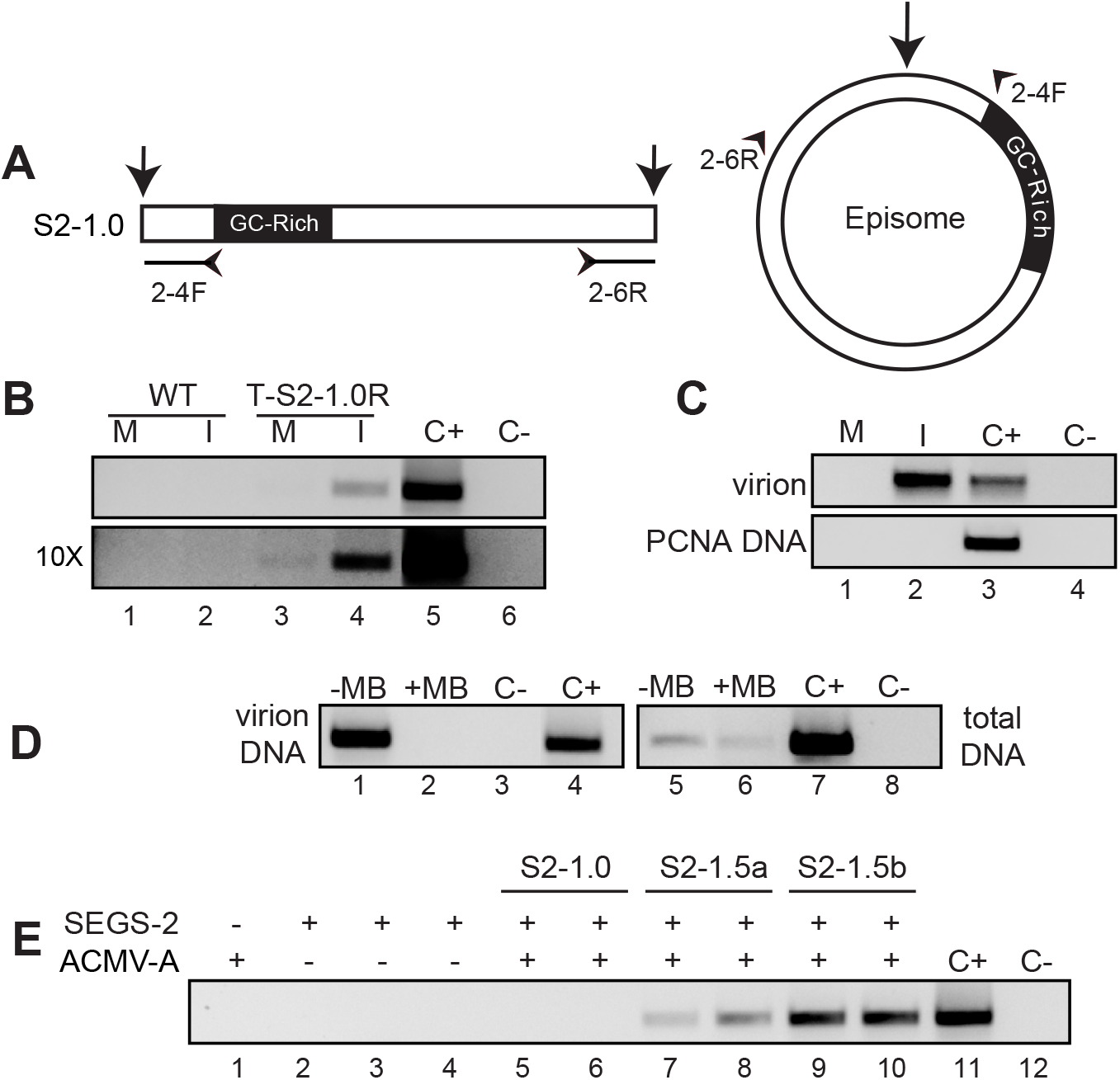
SEGS-2 episomes are packaged into virions. (A) The divergent primer pair 2-4F/6R (indicated by small arrowheads on the episome and linear model) was used to detect episomal copies after RCA of total DNA. The large arrows indicate the SEGS-2 episome junction site. (B) PCR products from mock (M) and ACMV-inoculated (I) wild-type (WT; lanes 1 and 2) and transgenic (T-S2-1.0R; lanes 3 and 4) Sei-0 leaves. The bottom gel, which is enhanced 10-fold, shows episomes in both mock and infected T-S2-1.0R leaves (lanes 3 and 4). (C) PCR of RCA products from virion DNA of mock-inoculated (M, lane 1) and ACMV-infected (I, lane 2) T-S2-1.0F plants. The convergent primer pair PCNA2-F/R verified that there was no genomic DNA contamination of virion RCA template (bottom gel lanes 1 and 2). (D) PCR of RCA products of virion and total DNA treated with Mung Bean nuclease (+MB, lanes 2 and 6) or not treated (-MB, lanes 1 and 5). (E) Amplification of SEGS-2 episome in replication assays using protoplasts from *N. tabacum* suspension cells. Episomes were analyzed in protoplasts co-transfected with ACMV DNA-A + S2-1.0 (lanes 5 and 6), S2-1.5a (lanes 7 and 8), or S2-1.5b (lanes 9 and 10). ACMV DNA-A was transfected alone in lane 1. The SEGS-2 plasmids were transfected alone in lanes 2 (S2-1.0), 3 (S2-1.5a), and 4 (S2-1.5b). The replication assays were repeated three times. The positive control (C+) used S2-1.5b plasmid DNA as template, and the negative control (C-) did not contain template DNA.

Geminiviruses and their satellites are packaged into virions as ssDNA, prompting us to ask if the SEGS-2 DNA in virions is single-stranded (Rojas et al., 2005). Virion and total DNA from ACMV-infected T-S2-1.0F plants were treated with Mung Bean nuclease, which only digests ssDNA. After treatment, virion and total DNA were used as templates in RCA reactions followed by PCR with divergent SEGS-2 primers. SEGS-2 episomal DNA was not detected in virion DNA after Mung Bean nuclease treatment (Fig. 3D, lane 2) even though episomes were observed in untreated virion DNA (lane 1). In contrast, SEGS-2 episomal DNA was observed in total DNA after Mung Bean digestion (Fig. 3D, lane 6), albeit at lower levels than for untreated total DNA (Fig. 3D, lane 5). These results established that SEGS-2 DNA is single-stranded in virions.

The observation that SEGS-2 episomes in total DNA samples are partially resistant to Mung Bean nuclease digestion is indicative of the existence of a double-stranded form. Geminivirus genomes replicate by a rolling circle replication mechanism that involves dsDNA (Hanley-Bowdoin et al., 2000). Hence, we asked if SEGS-2 also replicates in plant cells and if its replication is dependent on the presence of a helper virus. Protoplasts from *Nicotiana tabacum* suspension cells were transfected with the different SEGS-2 constructs in Fig. 1A either alone or in the presence ACMV DNA-A. Total DNA was extracted 48-h post transfection, treated with *Dpn*I to digest DAM-methylated input plasmid DNA, and analyzed by RCA followed by PCR with the primer pair 2-4F/6R to detect SEGS-2 episomes. SEGS-2 episomes were detected when S2-1.5a (Fig. 3E, lanes 7 and 8) or S2-1.5b (lanes 9 and 10) were cotransfected with the ACMV DNA-A replicon but not when ACMV DNA-A was absent (Fig. 3E, lanes 3 and 4). No SEGS-2 episomes were detected for S2-1.0 in the presence (Fig. 3E, lanes 5 and 6) or absence (lane 1) of ACMV DNA-A. These results established that SEGS-2 partial tandem dimer constructs, but not a monomer construct, support replication of a SEGS-2 episome in the presence of ACMV-A in *Nicotiana tabacum* suspension cells. More episomal SEGS-2 DNA was detected for S2-1.5b, indicating that it might contain two copies of the origin of replication (Stenger et al., 1991).

### The largest SEGS-2 ORF is necessary for its activity in *Arabidopsis thaliana*

SEGS-2 contains an ORF with the capacity to encode a 75 amino acid protein. To directly test if ORF 2 (Fig. 4A) is necessary for SEGS-2 activity, we mutated the ATG start codon in S2-1.5b. The activity of the ATG mutant was compared to wild-type SEGS-2 in Sei-0 plants co-bombarded with ACMV (Fig. 4B). The plants co-inoculated with ACMV and S2-1.5b began to show symptoms as early as 7 dpi (Fig. 4C) and went onto developed severe symptoms (Fig. 4B). In contrast, plants inoculated with ACMV alone or with ACMV and S2-ATGm began showing symptoms at 12 dpi (Fig. 4C). The plants coinoculated with ACMV and S2-ATGm fell into two group – one group (13 plants) that developed severe symptoms (S2-ATGmR) and another group (8 plants) that only showed mild symptoms (S2-ATGm) similar to plants inoculated with ACMV alone (Fig. 4B). DNA sequencing revealed that ATG mutation in the SEGS-2 ORF had reverted to wild-type in the plants with severe symptoms, while SEGS-2 could not be detected in plants with mild symptoms.

**FIG 4.**
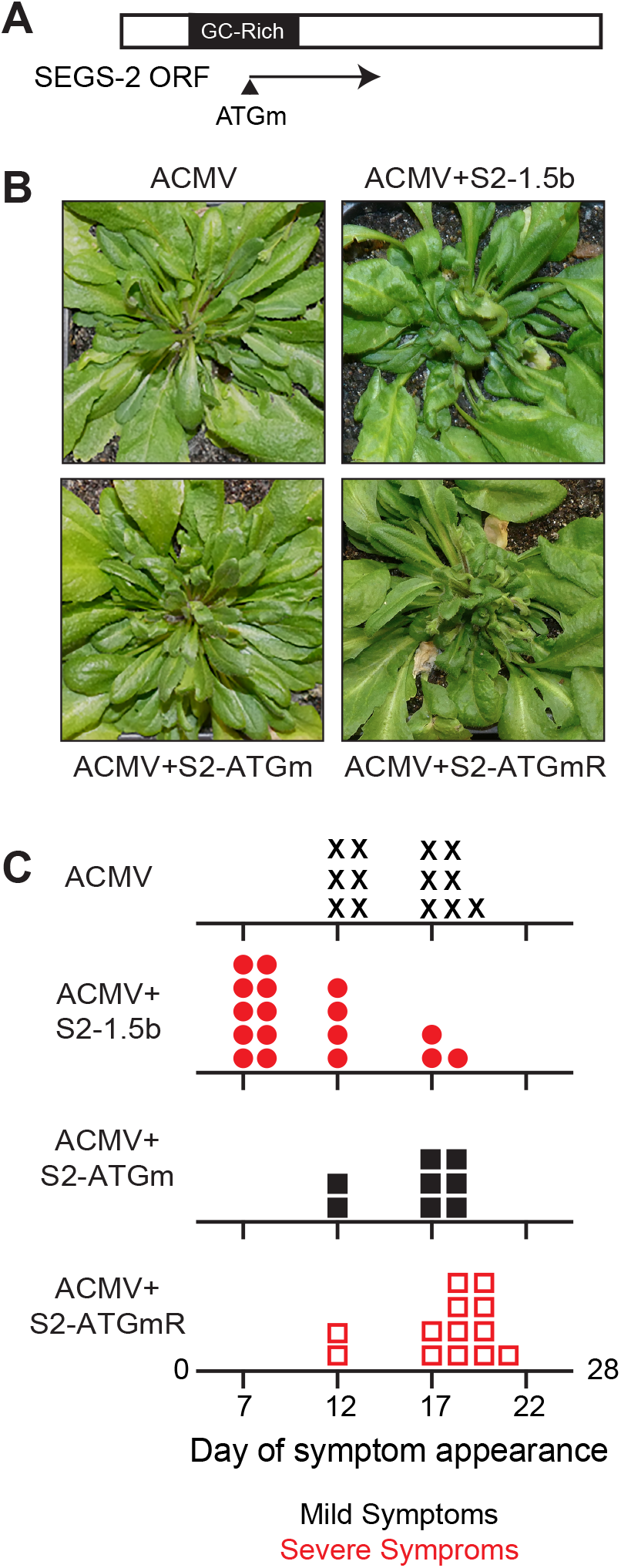
SEGS-2 encodes a functional open reading frame (ORF). (A) Diagram of the longest SEGS-2 ORF with mutated ATG indicated by arrow. (B) Arabidopsis Sei-0 plants inoculated with ACMV DNA-A and DNA-B alone or in combination with wild-type S2-1.5b or the ATG mutant (S2-ATGm) plasmid at 28 dpi. (C) Day of symptom appearance for individual plants with ACMV (X), ACMV + S2-1.5b (circle), ACMV + S2-ATGm (closed square), and ACMV + S2-ATGmRevertant (open square). Red indicates plants showing severe symptoms, and black indicates mild symptoms.

### Transcriptional analysis of SEGS-2 in transgenic lines

We next asked if SEGS-2 is transcribed in plants using RT-PCR and rapid amplification of cDNA ends (RACE). An 825-bp product was amplified from total RNA from healthy T-S2-1.0F and T-S2-1.0R plants by RT-PCR using the 2-hp1f/2-2BR primer pair (Fig. 5B). Detection of the product was independent of the orientation of the SEGS-2 insert in the T-DNA plasmid. The direction of transcription was determined using a 5’ RACE reaction. A 532-bp product (Fig. 5C, lane 2) matching the predicted size (positive control, lane 3) was observed using the LGSP1/2 primer pair in the LR (Left to Right) reaction. The band (Fig. 5C, lane 1) in the RR (Right Reaction) with the RGSP1/2 primers is of similar size to the band in the no template control (lane 4). Sequencing of the products from the two RACE reactions confirmed that the band in the LR reaction corresponds to SEGS-2 while the band in RR reaction is non-specific. Thus, transcription is orientated in the same direction as the SEG-2 ORF (Fig. 5A).

**FIG 5.**
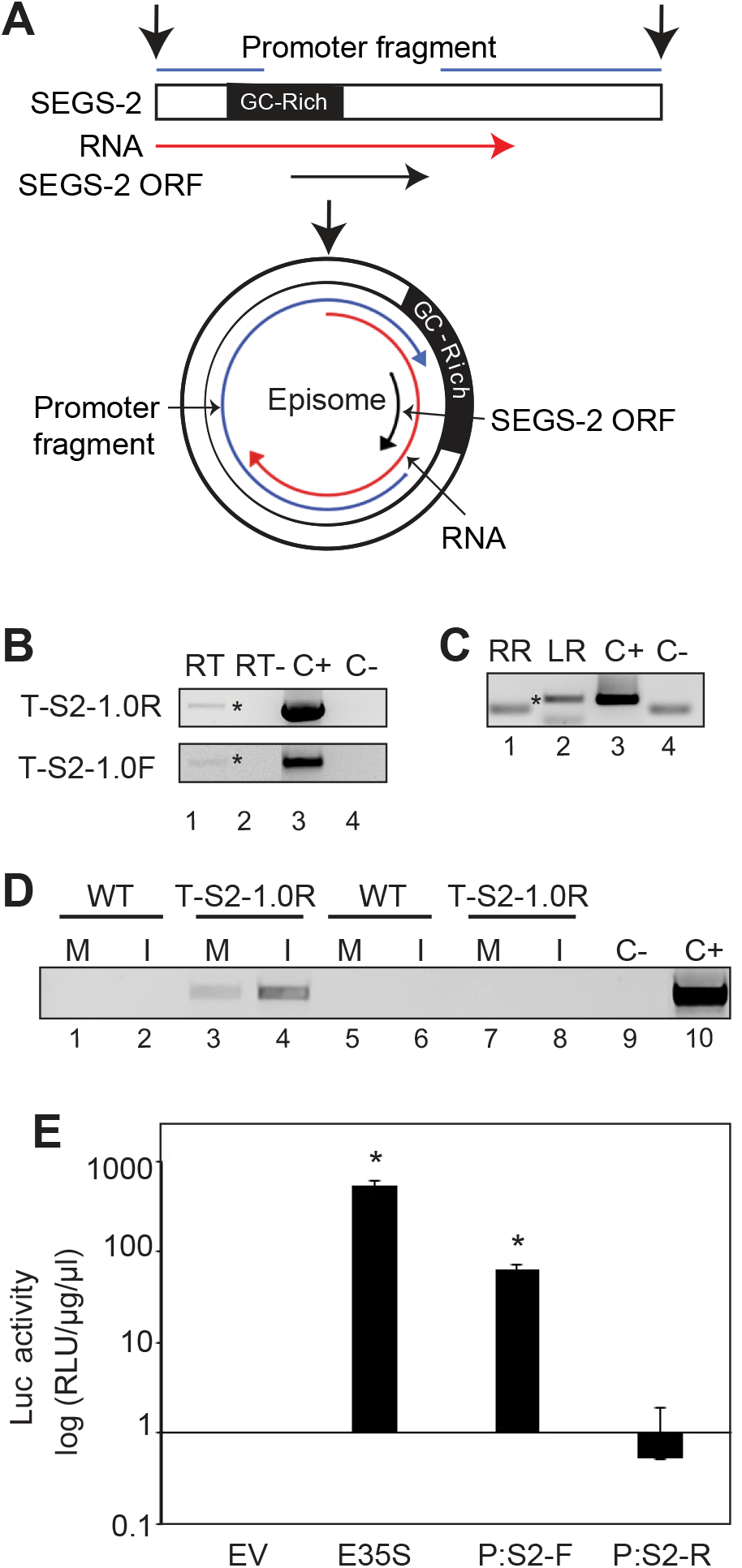
SEGS-2 transcription in healthy and infected plants. (A) Diagram of SEGS-2 transcription unit. Linear and episomal maps of SEGS-2 are shown, with the episomal junction marked by the vertical black arrows.The SEGS-2 ORF is marked by the horizontal and circular black arrows. The 825-bp SEGS-2 RNA is indicated by the red arrows The formation of the SEGS-2 episome brings a promoter (blue lines and arrow) in the 3’ end of linear SEGS-2 sequence to the 5’ end of the SEGS-2 ORF. (B) cDNA from total RNA extracted from healthy S2-1.0F and S2-1.0R (bottom) plants was amplified using convergent primer pairs, S2 hp1F/2BR and S2 7F/2R, respectively (RT, lane 1). The negative control did not include reverse transcriptase during cDNA synthesis (RT-, lane 2). (C) PCR of 5’ RACE reaction to the right (RR, lane 1) and left (LR, lane 2) with primer pair S2 hp1F/RGSP2. Asterisks in B and C indicate the SEGS-2 product. (D) RT-PCR (lanes 1-4) with primer pair S2 hp1F/2BR of cDNA from total RNA extracted (lanes 5-8) from wild-type Sei-0 (WT; lanes 1, 2, 5, and 6) and T-S2-1.0R (lanes 3,4,7 and 8) leaves in mock (M) and ACMV-inoculated (I) leaves. Lanes 5-8 are no reverse transcriptase negative controls. In B, C and D, the positive control (C+) used S2-1.0 plasmid DNA as template, and the negative control (C-) did not contain template. (E) Luciferase activity in tobacco protoplasts at 36 h after transfection with the forward (P:S2-F) and reverse orientation (P:S2-R) of the non-coding regions of SEGS-2 cloned upstream of the *Luc* gene. The E35S promoter and the empty vector (EV) are the positive and negative controls. Both P:S2-F and the E35S promoter displayed significant Luc activity compared to the EV control in a Students’ t-test (asterisks; P < 0.05). P:S2-R does not support *Luc* expression. Error bars correspond to two standard errors.

We then examined SEGS-2 transcripts during ACMV infection at 37 dpi using RT-PCR and the primer pair 2-hp1f/2-2BR. An 825-bp product was observed in total RNA from infected T-S2-1.0F plants (Fig. 5D, lane 4). More SEGS-2 RNA was detected in infected than mock inoculated T-S2-1.0F plants (Fig. 5D; cf. lanes 3 and 4). No SEGS-2 transcripts were detected in total RNA samples from mock-inoculated (Fig. 5D; lane 1) or infected (lane 2) wild-type plants.

The 5’ RACE product of the SEGS-2 transcript extended to the junction of the SEGS-2 insert and T-DNA vector. Given that detection of the SEGS-2 transcript was independent of the orientation of the SEGS-2 insert in the T-DNA (Fig. 5B), it was not clear what promoter sequences support SEGS-2 transcription. One possibility is that when SEGS-2 forms an episome, a sequence downstream of the transcript in the linear sequence functions as a promoter (Fig. 5A). To test this, we generated expression cassettes in which a 971-bp sequence from the stop codon to the 5’ ATG of the SEGS-2 ORF was inserted upstream of a luciferase reporter in both orientations (Fig. 5A). The cassettes were transfected into *N. tabacum* protoplasts and luciferase activity was measured 36 hours post-transfection (Fig. 4E). The sequence with the same orientation as the SEGS-2 ORF (P:S2-F) supported luciferase expression that was significantly above that of the empty vector control (Student T-test with p-value < 0.05). In contrast, the reverse orientation (P:S2-R) did not support luciferase expression. The amount of luciferase activity supported by the P:S2-F fragment was ca. 100-fold less than that of the strong enhanced 35S promoter (E35S), indicating that the SEGS-2 promoter is of moderate strength. Inclusion of ACMV DNA-A in the transfection reactions did not increase the activity of the SEGS-2 promoter (not shown).

### SEGS-2 enhances geminivirus infection in a resistant Arabidopsis accession

Pla-1 is an Arabidopsis accession that is resistant to *Cabbage Leaf Curl Virus* (CaLCuV) infection (Reyes et al., 2017). We asked if CaLCuV can infect Pla-1 plants when it is co-inoculated with SEGS-2. Pla-1 plants co-bombarded with CaLCuV and S2-1.5a or S2-1.5b displayed symptoms at 21 and 30 dpi (Fig. 6A) and showed no signs of flowering at 50 dpi (Fig. 6B). Consistent with the symptom results, viral DNA was detected at 29 and 43 dpi in plants co-inoculated with CaLCuV and SEGS-2 (Fig. 6C). In contrast, Pla-1 plants inoculated with CaLCuV alone resembled mock-inoculated plants in that they did not show symptoms or contain viral DNA (Fig. 6A, B, C). Immunochemistry analysis showed that the viral Rep protein was distributed throughout leaves from Pla-1 plants co-inoculated with CaLCuV and S2-1.5a but could not be detected in leaves from plants only inoculated with CaLCuV (Fig. 6D).

**FIG 6.**
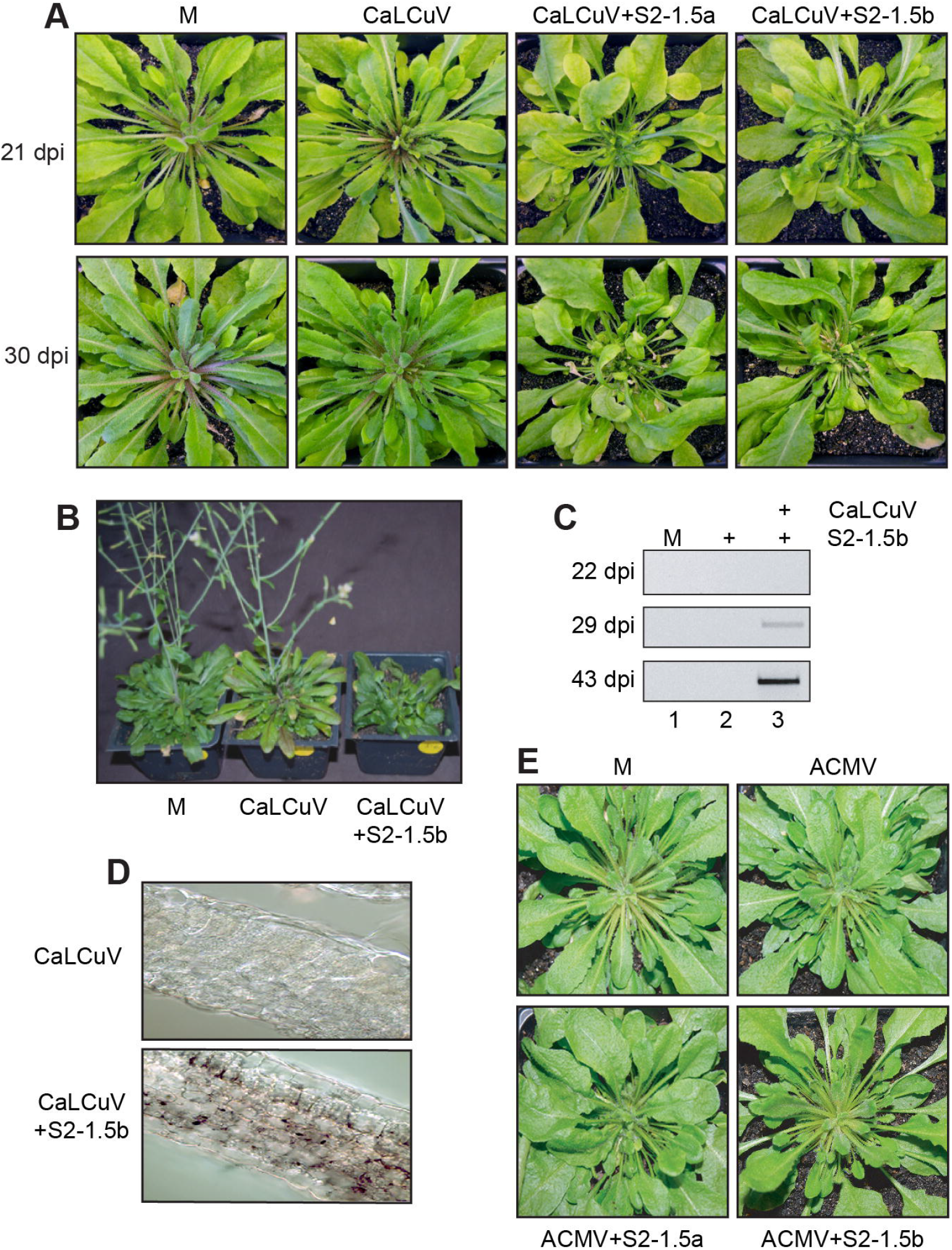
SEGS-2 facilitates CaLCuV infection of a resistant Arabidopsis accession. (A) Pla-1 plants inoculated with CaLCuV DNA-A and DNA-B alone or in combination with S2-2.5a or S2-1.5b plasmid at 21 and 30 dpi. (B) Flowering is suppressed in Pla-1 plants co-inoculated with CaLCuV and S2-1.5b but not with CaLCuV alone at 50 dpi. (C) Total DNA was extracted at 22, 29, and 43 dpi from systemically infected leaves and equivalent leaves from controls showing no symptoms, and analyzed by PCR for CaLCuV DNA-A with the divergent the primer pair CaLCuVFor/Rev. The lanes are mock inoculation (M, lane 1), CaLCuV + S2-1.5b (lane 2), or CaLCuV alone (lane 3). (D) Immunolocalization of CaLCuV Rep in Pla-1 leaf sections infected with CaLCuV alone or in combination with S2-1.5b at 36 dpi. CaLCuV Rep was only observed in leaves co-inoculated with CaLCuV and S2-1.5b. (E) Pla-1 plants inoculated with ACMV alone or in combination with S2-1.5a or S2-1.5b did not show symptoms at 30 dpi.

We also asked if ACMV can infect Pla-1 plants in the presence of SEGS-2. We failed to detect any symptoms when ACMV was cobombarded with S2-1.5a or S2-1.5b (Fig. 6E). Unlike CaLCuV, ACMV is not well-adapted to Arabidopsis and, as such, was not able to overcome resistance when it is co-inoculated with SEGS-2.

## DISCUSSION

Many begomovirus satellites enhance symptom severity and viral DNA accumulation (Briddon et al., 2003; Briddon and Stanley, 2006; Mansoor et al., 2003; Sivalingam and Varma, 2012; Zhou, 2013). In an earlier study, we showed that SEGS-2, a sequence that was amplified from CMD-infected cassava in Tanzania, also causes more severe symptoms during CMB infection of cassava (Ndunguru et al., 2016). SEGS-2 occurs as a circular episome in infected cassava and is packaged into virions, consistent with it being a DNA satellite. However, it has been difficult to study SEGS-2 in cassava because of the widespread presence of SEGS-2-related sequences in the cassava genome. To overcome this complication, we used heterologous plant systems to characterize SEGS-2 functionally in the absence of endogenous genomic sequences. We showed that like cassava, Arabidopsis plants infected with ACMV in the presence of SEGS-2 develop curling and filiform leaf symptoms. Symptom enhancement is dependent on maintenance of the largest ORF in SEGS-2. SEGS-2 also forms episomes that are transcribed and packaged as ssDNA into virions in Arabidopsis, and replicates in the presence of ACMV-A in tobacco cells. These properties strongly support our hypothesis that SEGS-2 is a new type of begomovirus satellite that contributes to increased CMD severity in Sub-Saharan Africa.

SEGS-2 increased ACMV symptoms in Arabidopsis when it was provided exogenously from a plasmid (Fig. 1) or from a transgene (Fig. 2). Plants co-inoculated with ACMV and the SEGS-2 monomer (S2-1.0) or partial tandem dimer clones (S2-1.5a or S2-1.5b) displayed stronger symptoms that appeared sooner compared to plants inoculated with ACMV alone. The configuration of the SEGS-2 clones had small but reproducible effects on symptoms later in infection, with S2-1.0 showing milder symptoms and a delay in appearance relative to S2-1.5a and S2-1.5b (Fig. 1D and 1E). The S2-1.5a and S2-1.5b clones were designed to resemble begomovirus partial tandem infectious clones, which are released from bacterial plasmid vectors via replication or recombination (Stenger et al., 1991). The same mechanisms could provide a rationale for why S2-1.5a and S2-1.5b show more symptom enhancement than S2-1.0.

Transgenic Arabidopsis carrying a SEGS-2 monomer sequence developed symptoms and accumulated viral DNA earlier and to higher levels than wild-type plants (Fig. 2B and 2D). These phenomena were independent of the orientation of the transgene indicating that T-DNA sequences did not impact SEGS-2 activity. Unexpectedly, T-S2-1.0 plants inoculated with ACMV alone displayed stronger symptoms than plants inoculated with exogenous S2-1.5b + ACMV (Fig. 2C). This may be due to the presence of the SEGS-2 transgene in every plant cell that is infected by the virus, which is unlikely when SEGS-2 is provided exogenously. Co-bombardment of S2-1.5b with ACMV in T-S2-1.0 plants did not increase symptoms compared to T-S2-1.0 plants inoculated only with ACMV, indicating that the transgene effect on symptoms was maximal and could not be further enhanced by the addition of exogenous SEGS-2 DNA (Fig. 2C). Together, these results support the idea that the availability of SEGS-2 throughout the plant is responsible for the stronger effect in transgenic Arabidopsis.

Earlier studies showed that the Pla-1 accession of Arabidopsis is immune to CaLCuV infection (Reyes et al., 2017). We showed that CaLCuV can infect Pla-1 plants in the presence of exogenous SEGS-2 (Fig. 6). This result was unexpected because studies in cassava indicated that SEGS-2 cannot overcome CMD2 resistance to CMBs (Ndunguru et al., 2016). This difference may reflect dissimilarities in the resistance mechanisms, which are not yet known for either Arabidopsis or cassava. The ability of SEGS-2 to assist CaLCuV infection in overcoming resistance indicated that its activity is not specific for CMB infection, and instead, is likely an effect on a general feature of the begomovirus infection process. An interesting feature of CaLCuV infection in Pla-1 is the presence of virus throughout the leaf in the presence of SEGS-2 (Fig. 6D). In the susceptible accession Col-0, CaLCuV is vascular associated, and is not commonly seen in mesophyll or epidermal cells (Ascencio-Ibáñez et al., 2008). It is not known if the lack of detectable virus in Pla-1 plants in the absence of SEGS-2 is due to inhibition of viral replication or movement of virus out of the inoculated cell. The widespread distribution of CaLCuV in the presence of SEGS-2 in Pla-1 suggest that SEGS-2 breaks resistance by overcoming a barrier to viral cell-to-cell movement and the establishment of infection. It is noteworthy that ACMV cannot infect Pla-1 in the presence of SEGS-2 (Fig. 6E), suggesting that the barrier to infection is greater for a virus poorly adapted to Arabidopsis.

An earlier study showed that SEGS-2 episomes are encapsidated into virions in CMB-infected cassava plants and viliferous whiteflies (Ndunguru et al., 2016). ACMV-infected Arabidopsis with a SEGS-2 transgene also contained SEGS-2 episomes (Fig. 3B) that were packaged into virions (Fig. 3C). We extended these observations by showing that SEGS-2 is packaged as ssDNA in Arabidopsis. At 1.2 Kb, SEGS-2 is similar in size to other begomovirus satellites, which are also encapsidated into virions as ssDNA (Hesketh et al., 2018; Saunders et al., 2020). SEGS-2 also occurs as a dsDNA in Arabidopsis leaves, characteristic of rolling circle and/or recombination dependent replication (Hanley-Bowdoin et al., 2013; Jeske, Lütgemeier, and Preiss, 2001) (Fig. 3D). Replication assays in tobacco protoplasts established that SEGS-2 can replicate in the presence of ACMV-A (Fig. 3E) analogous to betasatellites, which also rely on begomovirus helper viruses for their replication (Xu et al., 2019; Zhou, 2013). DNA minicircles, which consist of both viral and host DNA, have been identified in begomovirus infected plants (Catoni et al., 2018). Like SEGS-2, the minicircles, which are packaged into virions, can also replicate in the presence of a helper begomovirus. However, they differ from SEGS-2 in that they contain viral DNA with an origin of replication, while SEGS-2 does not contain a recognizable viral replication origin.

The replication properties of the different SEGS-2 clones may provide some insight into the origin of replication in SEGS-2. S2-1.5a and S2-1.5b replicated in the presence of ACMV-A, but no replication was detected for S2-1.0. In addition, S2-1.5b replicated to higher levels than S2-1.5a. S2-1.5a and S2-1.5b contain a 21-bp inverted repeat with a 4-bp loop that includes the SEGS-2 episomal junction. The 3’ stem sequence, which does not occur in the cassava genome, shows strong similarity to alphasatellite origin sequences (Ndunguru et al., 2016). S2-1.0 does not include the stem-loop because the 21-bp sequences are separated by intervening SEGS-2 sequences in the cloned DNA. If the hairpin is involved in SEGS-2 replication, this would provide an explanation for why S2-1.0 did not replicate in the transient assays. The difference between S2-1.5a and S2-1.5b may reflect that different regions of SEGS-2 are duplicated in the partial tandem dimers. If S2-1.5b contains two functional origins flanking SEGS-2 sequences, this configuration would facilitate replicational release of an episome. In contrast, without two functional origins, S2-1.5a could release a SEGS-2 episome via recombination between the duplicated sequences, a process that is less efficient than replicational release (Stenger et al., 1991). The end result would be the detection of more nascent SEGS-2 DNA for S2-1.5b than for S2-1.5a in the replication assays.

The observation that the S2-1.0 clone does not replicate in transient replication assays raises the question as to why the SEGS-2 monomer forms episomes during infection in Arabidopsis. Although the 21-bp sequences at the ends of the SEGS-2 insert in S2-1.0 do not facilitate episome formation during the 48-h replication assays, they may enable episome formation over longer periods of time in transgenic T-S2-1.0 plants. The detection of SEGS-2 episomes at very low levels in mock-inoculated T-S2-1.0 plants established that SEGS-2 episome formation is not dependent on virus infection and, instead, is a property inherent in the SEGS-2 monomer sequence. SEGS-2 episomes could have been generated at any time during growth of T-S2-1.0 plants or in response to wounding during bombardment (Cheong et al., 2002; Marhava et al., 2019). A change in DNA methylation status due to wounding may trigger the release of the SEGS-2 monomer at the site of the inverted repeats and the formation of an episome (Colicchio, Kelly, and Hileman, 2020). The presence of SEGS-2 episomes in mock-inoculated plants also provides an explanation for the detection of SEGS-2 RNA in the mock controls. Once generated, SEGS-2 episomes can undergo replication during ACMV infection, resulting in the observed increase for both episomes and SEGS-2 RNA in infected plants.

Like betasatellites, SEGS-2 encodes a small protein that is necessary for its activity in Arabidopsis (Xu et al., 2019; Zhou, 2013). Analysis of SEGS-2 ATG mutants uncovered two ACMV infection phenotypes – one with mild symptoms and one with severe symptoms (Fig. 4B). The timing of symptom appearance was delayed for the severe symptom class compared to the wild-type SEGS-2 control (Fig. 4C). Sequencing revealed that the SEGS-2 mutation had undergone reversion restoring the SEGS-2 ORF in all of the plants with severe symptoms. High levels of reversion have also been reported for *AC1* mutants in Tomato golden mosaic virus (Arguello-Astorga et al., 2007). In that case, the mutant *AC1* virus reverted in all infected plants within 3 weeks, reflecting strong selection pressure to restore interactions between the viral Rep protein and a host protein necessary for efficient viral replication (Kong et al., 2000). Reversion of the SEGS-2 ORF did not occur in all plants, most likely because, unlike Rep, the SEGS-2 protein is not essential for begomovirus infection. However, the high level of reversion supports the importance of the SEGS-2 ORF during infection of a poorly adapted virus like ACMV in Arabidopsis.

The SEGS-2 ORF specifies a protein containing 75 amino acids. The cassava genome contains two small ORFs with the potential to encode 69 amino acid proteins with high similarity to SEGS-2. The small ORFs are located within a 38-Kb region on chromosome 13 that includes the previously identified SEGS-2 partial copies, PC2-2 and PC2-3 (Ndunguru et al., 2016). The PC2-2 and PC2-3 ORFs show 85% and 89% amino acid identity to SEGS-2 ORF, respectively (Ndunguru et al., 2016). The PC2-2 ORF is not associated with a cassava gene, and there is no evidence that it is transcribed. In contrast, the PC2-3 ORF is located in the 5’ UTR of an mRNA transcribed from a cassava gene encoding a PPR repeat protein (Manes.13G072800). Small ORFs in 5’ UTRs have been implicated in negative control of translation, and they can impact protein expression in a variety of plant processes, including the host defense response (Zhang et al., 2020). SEGS-2 -related sequences in the cassava genome are often found in 5’ UTRs, suggesting that they may be involved in translational control of many genes. However, it is not known if the proteins specified by PC2-2 and PC2-3 have roles other than translational control, analogous to the SEGS-2 protein (Ndunguru et al., 2016).

Our results support a model in which SEGS-2 was generated by a recombination event between a cassava genomic sequence and an alphasatellite (Fig. 7). The recombinant DNA forms an episome that is enscapsidated into virions as ssDNA. SEGS-2 also exists as a dsDNA that is formed during replication in the presence of a helper virus. SEGS-2 encodes an ORF that is transcribed from the viral dsDNA template. The SEGS-2 ORF specifies a small protein that enhances symptoms, potentially by altering cell-to-cell movement of viral DNA. Taken together, we propose that SEGS-2 represents a new type of low-copy number begomovirus satellite with the capacity to enhance disease and break resistance. Future studies that focus on the interactions of SEGS-2 with CMBs and cassava may contribute to the development of effective and sustainable disease control measures for CMD.

**FIG 7.**
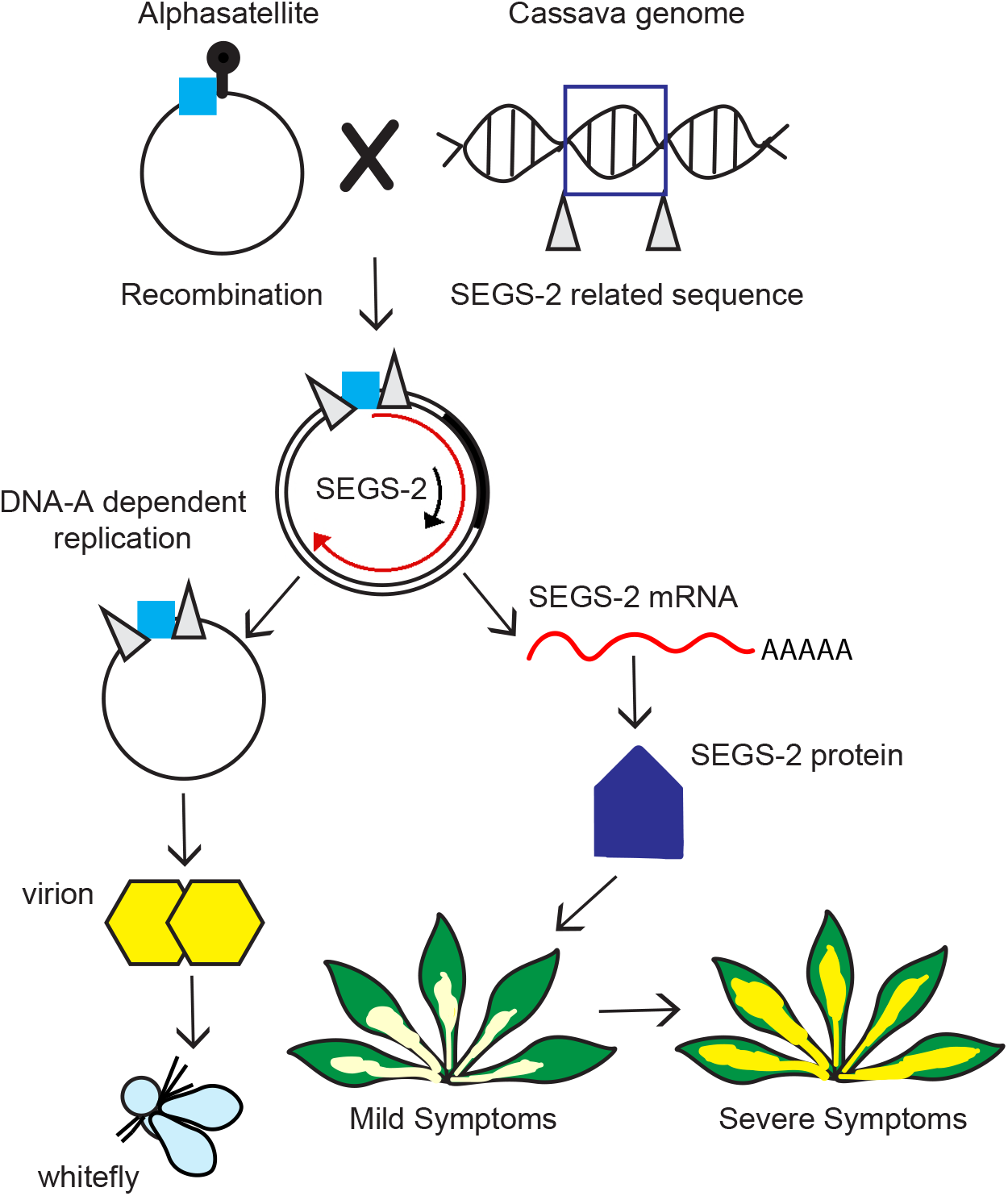
Model of SEGS-2 biology. A SEGS-2 sequence in the cassava genome recombined with an alphasatellite and acquired a small fragment adjacent to the satellite origin of replication (blue box). The alphasatellite sequence may facilitate rolling circle replication of a circular SEGS-2 episome in the presence of a CMB helper virus. The SEGS-2 episome is packaged into virions and transmitted by whiteflies as part of the CMB virus complex. The SEGS-2 episome is transcribed and the RNA is translated to produce a 75-amino acid protein that is necessary for the enhancement of geminivirus symptoms.

## MATERIAL AND METHODS

### Cloning

All of the plasmids and primers used in this study are listed in Table 1 and Table S1. All of the clones and mutants were confirmed by Sanger sequencing.

**Table 1.**
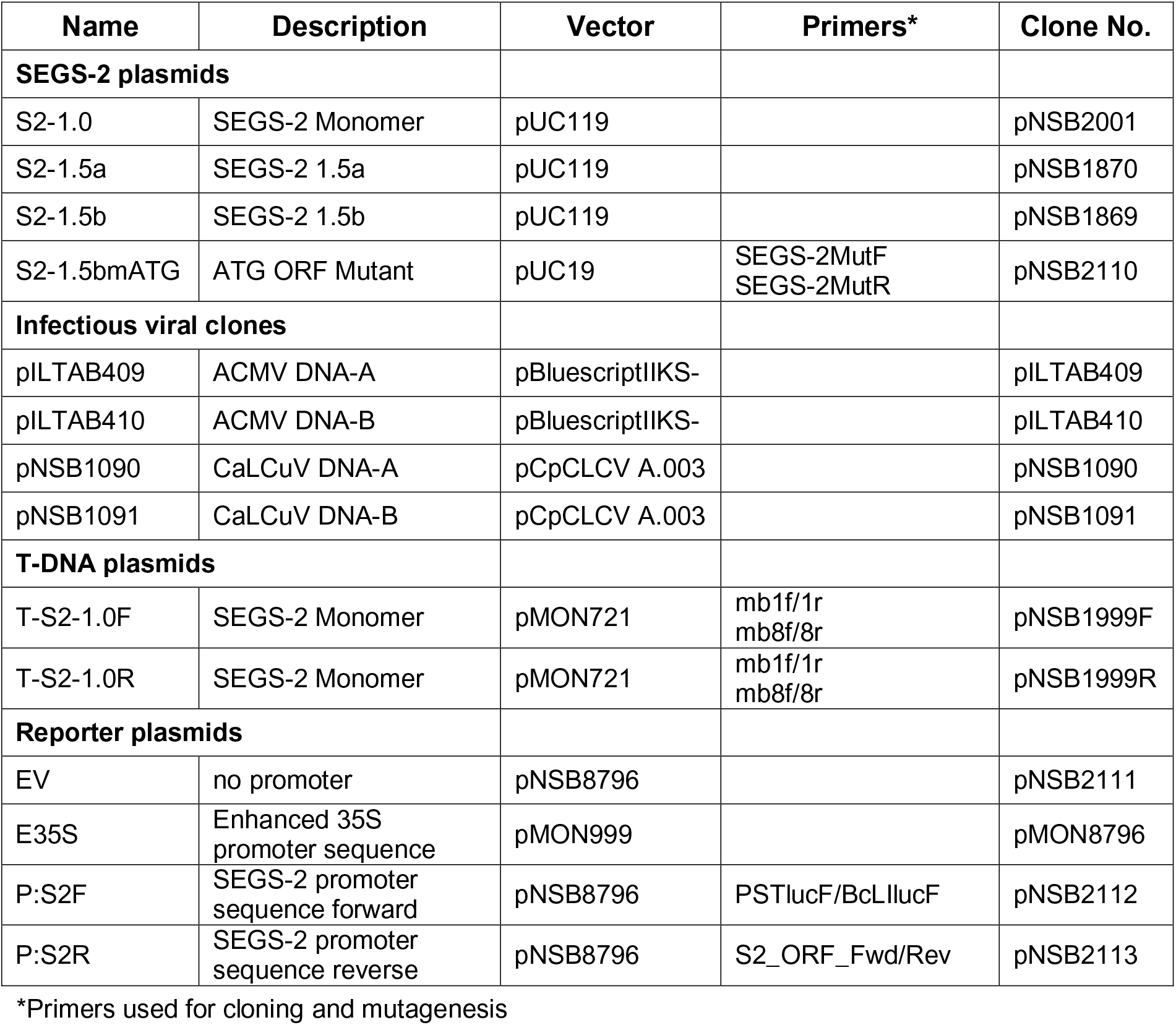
Plasmids used for plant studies.

| Name | Description | Vector | Primers* | Clone No. |
| --- | --- | --- | --- | --- |
| **SEGS-2 plasmids** |  |  |  |  |
| S2-1.0 | SEGS-2 Monomer | pUC119 |  | pNSB2001 |
| S2-1.5a | SEGS-2 1.5a | pUC119 |  | pNSB1870 |
| S2-1.5b | SEGS-2 1.5b | pUC119 |  | pNSB1869 |
| S2-1.5bmATG | ATG ORF Mutant | pUC19 | SEGS-2MutF<br>SEGS-2MutR | pNSB2110 |
| **Infectious viral clones** |  |  |  |  |
| pILTAB409 | ACMV DNA-A | pBluescriptIIKS- |  | pILTAB409 |
| pILTAB410 | ACMV DNA-B | pBluescriptIIKS- |  | pILTAB410 |
| pNSB1090 | CaLCuV DNA-A | pCpCLCV A.003 |  | pNSB1090 |
| pNSB1091 | CaLCuV DNA-B | pCpCLCV A.003 |  | pNSB1091 |
| **T-DNA plasmids** |  |  |  |  |
| T-S2-1.0F | SEGS-2 Monomer | pMON721 | mb1f/1r<br>mb8f/8r | pNSB1999F |
| T-S2-1.0R | SEGS-2 Monomer | pMON721 | mb1f/1r<br>mb8f/8r | pNSB1999R |
| **Reporter plasmids** |  |  |  |  |
| EV | no promoter | pNSB8796 |  | pNSB2111 |
| E35S | Enhanced 35S promoter sequence | pMON999 |  | pMON8796 |
| P:S2F | SEGS-2 promoter sequence forward | pNSB8796 | PSTlucF/BcLlucF | pNSB2112 |
| P:S2R | SEGS-2 promoter sequence reverse | pNSB8796 | S2_ORF_Fwd/Rev | pNSB2113 |

\*Primers used for cloning and mutagenesis

The SEGS-2 clones were generated from a pGEM-T Easy plasmid harboring a dimeric copy of SEGS-2 (pGEM-2SEGS-2) (Ndunguru *et al.,* 2016). pGEN-2SEGS-2 was digested with *Eco*R1 to release the dimer and ligated into pUC119 also cut with *Eco*RI to create pNSB2137. A 1197-bp SEGS-2 sequence in pNSB2137 was released using *Bam*HI and ligated into pUC119 linearized with *Bam*HI to generate the monomer clone, S2-1.0 (pNSB2001). S2-1.5a (pNSB1870) and S2-1.5b (pNSB1869) are partial tandem dimer clones with one full copy of SEGS-2 and duplicated sequences corresponding to SEGS-2 positions 1-449 and 449-1197, respectively. A 586-bp SEGS-2 sequence was released from pNSB2137 as a *Nde*I-repaired/*Nsi*I fragment and ligated into pUC119 digested with *Pst*I and *Sma*I to generate pNSB1868. A 748-bp fragment was released from pNSB2001 as a *Bam*H1/*Nde*I-repaired fragment and ligated into pUC119 digested with *Bam*HI and *Sma*I to generate pNSB1867. pNSB2001 was digested with *Bam*HI to release a 1197-bp SEGS-2 fragment and ligated into both pNSB1868 and pNSB1867 also digested with *Bam*HI to give S2-1.5a (pNSB1870) and S2-1.5b (pNSB1869), respectively. The SEGS-2 ATG mutant (pNSB2110) was created in S2-1.5b (pNSB1869) using the QuickChange II Site-Directed Mutagenesis Kit (Agilent Technologies) and the primers, SEGS-2MutF/SEGS-2MutR.

*Not*I sites were introduced at the ends of the S2-1.0 insert in pNSB2001 using the QuikChange II Site-Directed Mutagenesis Kit and the primers, mb1f/1r-mb8f/8r (Table S1). The resulting fragment was digested with *Not*I to release the SEGS-2 fragment, which was cloned into the T-DNA plasmid pMON721 linearized with *Not*I to give T-S2-1.0 (pNSB1999F/R). The orientation of the SEGS-2 insert in the vector was determined by *Pst*I digestion. *Agrobacterium tumefaciens* strain ABI was transformed via electroporation with both the forward (T-S2-1.0F) and reverse (T-S2-1.0R) orientations (Shaw, 1995) (Table S1).

The P:S2-F luciferase reporter cassette (pNSB1949) was generated from pMON1869 using the primers, PST_Promoter_Luciferase/BCII_Promoter_Luciferase (Table S1). The resulting 973-bp fragment was digested with *Pst*I and *Bcl*I and ligated upstream of the luciferase coding sequence in pMON8796 that was first digested with *Pst*I and *Bcl*I to remove the E35S promoter. The P:S2-R cassette was inserted into pMON8796 using the NEB Builder Kit with primer pair S2-ORF_Fwd/Rev to amplify the 973-bp cassette. pMON9708empty_Fwd/Rev was used to amplify the pMON8796 vector with matching overlapping ends to the P:S2-R cassette following NEB Builder Kit protocol (See Table S1).

### Arabidopsis Transformation

The *Arabidopsis thaliana* accession Sei-0 from the Arabidopsis biological resource center (ABRC, Ohio State University: Stock Number CS1504) was transformed using the floral dip method (Mara, Grigorova, and Liu, 2010) to introduce T-S2-1.0F and T-S2-1.0R. Seeds of Arabidopsis transformants were sterilized in 1.5 mL Eppendorf tubes with 70% ethanol for 2 min, followed by treatment with a 5% bleach solution containing 0.02% Triton X-100 for 10 min and washing 3 times with sterile water. Approximately 100 sterile seeds were placed on selection plates (MSO medium with 0.5 mg/mL kanamycin) and sealed with aeration tape. The plates were placed in the dark at 4°C for 2 days followed by a brief exposure to light for 6 h at room temperature and incubated again for 2 days in the dark at 4°C until the seeds began to germinate. The plates were then placed in constant light at room temperature for plantlet growth. Three sequential generations were screened for kanamycin resistance. Second generation (T2) of seeds that showed 3:1 segregation was screened to identify homozygous lines in the third generation (T3).

### Infection Assays

Plants corresponding to the Arabidopsis accessions, Sei-0, Col-0 and Pla-1, were grown at 20°C under 8-h light/16-h dark cycles. For infection, 6 to 7-week-old plants with ca. 30 leaves were inoculated using a microsprayer at 30 psi to deliver gold particles coated with SEGS-2 plasmids (600 ng/6 plants) alone or in combination with infectious clones for ACMV (5 µg each of DNA-A and DNA-B/ 6 plants; Accession Numbers: MT858793.1 and MT858794.1 (Hoyer et al., 2020)) or CaLCuV (2.5 µg each of DNA-A and DNA-B/ 6 plants; Accession Numbers: NC_003866 and NC_003887.1 (Abouzid A. M., 1992)). Nine plants were inoculated for each DNA combination treatment, and a mock inoculation control was included for each experiment. All experiments were repeated three times. Disease symptoms were monitored visually starting at 7 dpi and continuing for up to 35 dpi. Symptoms, including chlorosis, leaf deformation and stunting, were scored on 1-5 symptom severity scale (scale: 1 = no symptoms to 5 = very severe) throughout new growth in the rosette. Total DNA was isolated from Arabidopsis plants at 7, 14, 21, 28, and 35 dpi using the CTAB protocol (Doyle JJ, 1990). ACMV or CaLCuV DNA-A accumulation was characterized in total DNA (0.1μg) samples by semi-quantitative PCR using the primer pairs, CMAFor2/CMARev2 and CaLCuV-For/CaLCuV-Rev, respectively. Both primer sets consisted of divergent primers that only amplified circular DNA-A molecules released from infectious clone DNA efficiently. For CaLCuV, PCR using Standard Taq Polymerase (NEB) was performed for 30 cycles of (denaturation: 45 s at 95°C, annealing: 45 s at 53°C; extension: 2.25 min at 72°C). PCR conditions for ACMV were similar except the annealing temperature was 55°C and Hot Start Taq Polymerase (NEB optimized for large product sizes was used. The PCR products (885 bp for CaLCuV and 2593 bp for ACMV) were resolved on 1% (w/v) agarose gels and stained with ethidium bromide for UV light visualization.

### Analysis of SEGS-2 RNA

For transcript analysis, total RNA was isolated from CMB-infected and mock-inoculated leaves at 35 dpi using the mirVana^TM^ miRNA Isolation Kit (Ambion, Carlsbad). Total RNA (10 μg) was subjected to DNase I digestion using DNA-free^TM^ kit (Ambion), and DNase-treated RNA (1 μg) was used for first-strand cDNA synthesis by M-MuLV reverse transcriptase (200 U for 1 h at 42°C). The transcript was amplified using the primer pair, 2-hp1F/2-2BR. PCR, for 37 cycles (1 min at 95°C, 1 min at 49°C, 1 min at 72°C). The product (825 bp) was resolved on 1% (w/v) agarose gels, stained with ethidium bromide for UV light visualization, and sequenced.

The 5’ end of the transcript was determined using Rapid Amplification of cDNA Ends (RACE) (Yeku and Frohman, 2011). Total RNA (1 μg) from healthy Arabidopsis leaves was heated to 75°C for 5 min followed by reverse transcription with M-MuLV reverse transcriptase (200 U for 1 hour at 42°C) with a left or right gene-specific primer (LGSP1 or RGSP1, Table S1). The cDNA synthesis product was purified using QIAquick PCR Purification Kit (Qiagen, Maryland), and used in a dA-tailing reaction with terminal transferase (120 U for 1.5 h at 37°C). Amplification of the target cDNA was performed using a second set of left (LGSP2) or right (RGSP2) gene-specific primers with Adaptor(dT)17 that binds to the poly(A)^+^ tract added in dA-tailing reaction and the Adaptor primer that binds to the 5’ end of the Adaptor(dT)17. PCR was performed for RGSP2 for 1 cycle (5 min at 94°C, 5 min at 54°C, 40 min at 72°C), followed by 30 cycles (40 s at 94°C, 1 min at 54°C, 3 min at 72°C) and a final cycle (40 s at 94°C, 1 min at 54°C, 15 min at 72°C). PCR cycle conditions for LGSP2 were similar except the annealing temperature was 52°C. The PCR products for primer pairs, RGSP2 (257 bp) and LGSP2 (546 bp), were gel purified using QIAquick Gel Extraction kit (Qiagen) and sequenced using the RGSP2 or LGSP2 primer, respectively.

### Analysis of SEGS-2 episomes in *Arabidopsis thaliana*

Total DNA was isolated from 1 mg of pooled Arabidopsis leaf 3 tissue collected from three plants for each treatment. Virion samples were generated by homogenizing tissue collected from 3 infected leaves (1 mg) in 50 mM Tris, 10 mM MgSO_4_, 0.1 M NaCl, pH 7.5, followed by low speed centrifugation. The supernatant was subjected to 0.22 μM filtration followed by DNase I digestion (2.5 U for 3 h at 37°C). Virion DNA was isolated using QIAamp MinElute Virus Spin Kit (Qiagen, Maryland).

Total and virion DNA were amplified by rolling circle amplification (RCA) using the templiPhi100 DNA Amplification Kit (GE Healthcare) according to the manufacturer’s instruction. The RCA product was diluted 10-fold with DNase-free water and 1 μ was used as template in a 50-μL PCR reaction containing the primer pair, S2-4F and S2-6R (Table S.1), using previously established conditions (Ndunguru et al., 2016). The RCA products were tested for Arabidopsis genomic DNA contamination using the primer pair, PCNA2-F/PCNA2-R, according to (Ndunguru et al., 2016).

To determine if the SEGS-2 episomes were single- or double-stranded DNA, total or virion DNA was treated with 1 unit of Mung Bean nuclease (NEB, Massachusetts). The digestion products were analyzed for SEGS-2 episomes as described above.

### Luciferase and *replication* assays

SEGS-2 promoter assays in *N. tabacum* NT-1 cell protoplasts were performed according to published protocols (Eagle et al., 1994). The luciferase reporter cassettes (10 μg DNA/2.8 × 106 cells) corresponded to the Empty Vector (no promoter, negative control), the enhanced 35S promoter (positive control) (Eagle et al., 1994), and the P:S2-F and P:S2-R cassettes. The protoplasts were cultured for 36 h after transfection, resuspended in 250 μL of extraction buffer (100 mM KH_2_PO_4_, 1 mM EDTA, 10 mM DTT, 8 mM phenylmethylsulfonyl fluoride, 0.5% glycerol, pH 7.8), sonicated and centrifuged. Luciferase activity was measured in the resulting supernatants as previously described (Eagle et al., 1994). The assays were repeated four times and a two-tailed Student’s t-test was used to compare mean *luc* activities.

For SEGS-2 replication assays, NT1 protoplasts were electroporated with SEGS-2 plasmids (10 μg/2.8 × 10^6^ cells) in the presence or absence of an ACMV DNA-A replicon (1.5 μg/2.8 × 10^6^ cells) and cultured as described previously (Fontes et al., 1994). Total DNA was purified 48 h post transfection, and 30 μg was digested with *Dpn*I. Episomal forms of SEGS-2 were analyzed using divergent primer pairs 2-4F/2-6R as described above.

### *In situ* hybridization and immunohistochemical staining

Leaves 5 and 6 relative to the center of the rosette were harvested at 35 or 36 dpi from S2-1.0F or Pla-1 plants, respectively, fixed using paraformaldehyde, and embedded in 5% low-melting point agarose in 1xPBS buffer as described previously (Shen and Hanley-Bowdoin, 2006). The leaf tissue was cut into 100-μm sections using a Leica VT1000S vibratome (Leica Microsystems).

For *in situ* hybridization, a digoxigenin-labeled probe corresponding to 415 bp of the ACMV AC1 gene was generated using a PCR DIG Probe Synthesis kit (Roche Diagnostics) and the primer pair, ACMV 400F and ACMV 400R (Table S1). PCR was performed in a 50 μL reaction mix containing 10 ng of ACMV-A DNA with the primer pair above (Table S1) according to the manufacturer’s instructions. Labeled and unlabeled PCR products were analyzed on 1% agarose gels to determine labeling success. Before use, the probe was denatured at 100°C for 5 min and cooled on ice for 5 min.

Immunohistochemistry used a rabbit antiserum against CaLCuV Rep, a biotinylated anti-rabbit IgG secondary antibody and horse radish peroxidase conjugated to streptavidin. Preparation procedures, the source of the antibodies and immunohistochemistry protocol were described previously (Ascencio-Ibáñez et al., 2008; Shen et al., 2014).

## ACKNOWLEDGEMENTS

We thank Dr. Wei Shen for his help in manuscript review and Dr. Vincent Fondong for the ACMV clones. This study was funded by grants from the National Science Foundation (DBI-1110050) to LH-B and JN and the Bill and Melinda Gates Foundation (OPP1149990) to LH-B, JTA-I and JN. CDA was supported with NSF Graduate Research Fellowship and an NCSU Provost Fellowship. LDLG was supported by a Fulbright Fellowship.

## Supporting information

**S1 Table 1.**
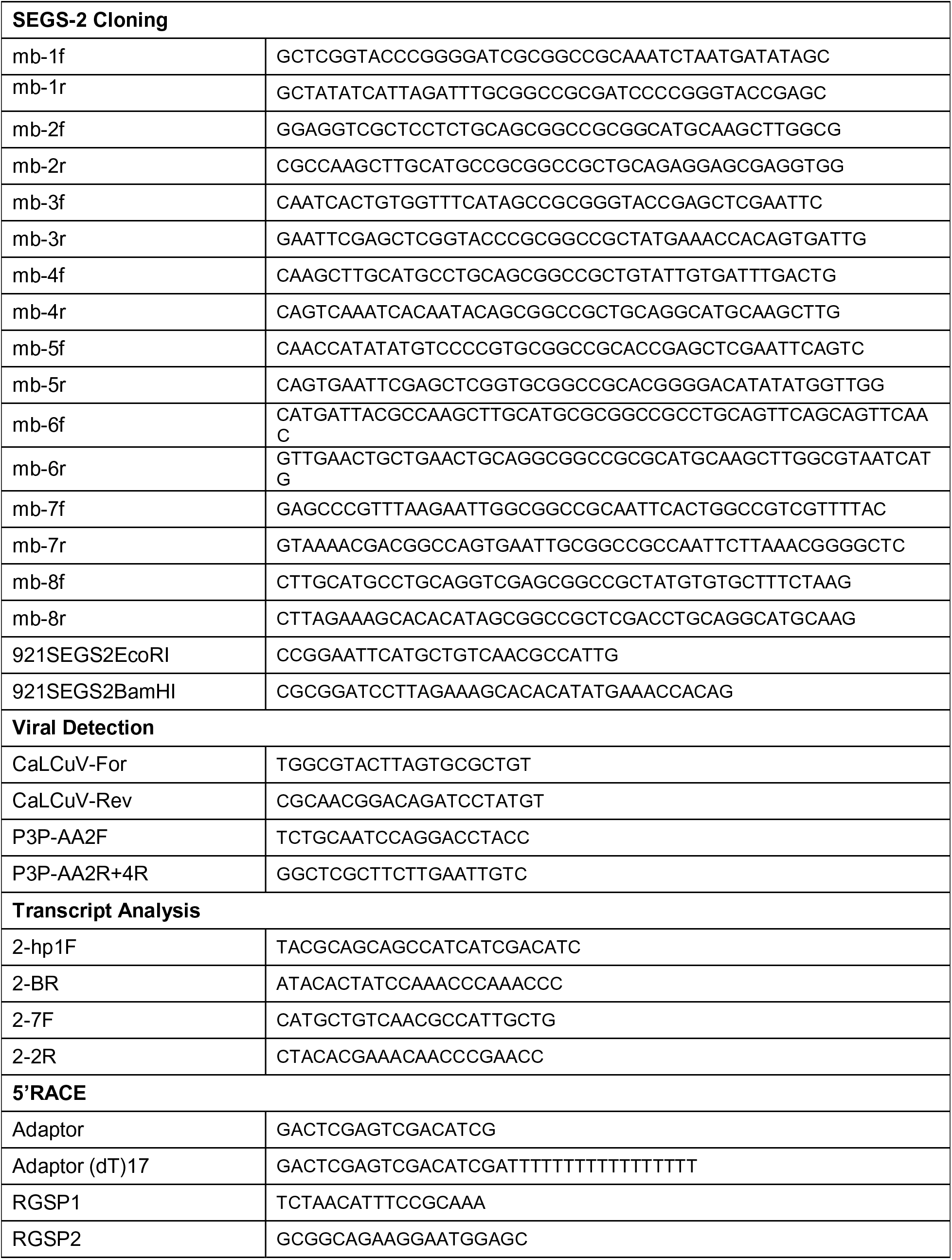
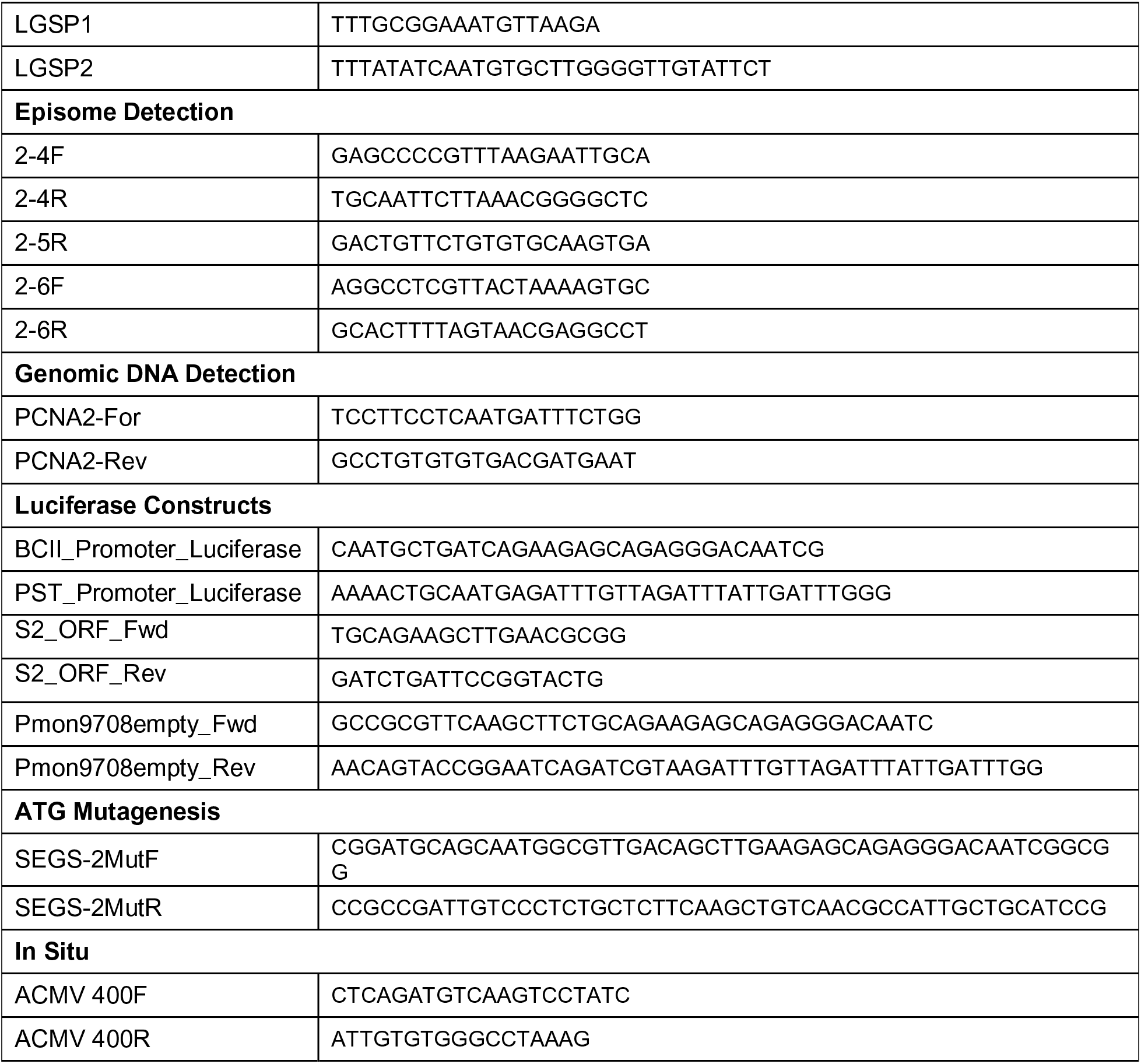
PCR Primer Sequences used in this study.

## Notes

### Competing Interest Statement

The authors have declared no competing interest.

